# Comparison between O and O_H_ intermediates of cytochrome *c* oxidase studied by FTIR spectroscopy

**DOI:** 10.1101/275099

**Authors:** E. Gorbikova, R. Kalendar

## Abstract

Cytochrome *c* oxidase is terminal enzyme in the respiratory chain of mitochondria and many aerobic bacteria. It catalyzes reduction of oxygen to water. During its catalysis, C*c*O proceeds through several quite stable intermediates (**R**, **A**, **P**_**R/M**_, **O/O**_**H**_, **E/E**_**H**_). This work is concentrated on the elucidation of the differences between structures of oxidized intermediates **O** and **O**_**H**_ in different C*c*O variants and at different pH values. Oxidized intermediates of wild type and mutated C*c*O from *Paracoccus denitrificans* were studied by means of static and time-resolved Fourier-transform infrared spectroscopy in acidic and alkaline conditions in the infrared region 1800-1000 cm^-1^. No reasonable differences were found between all variants in these conditions, and in this spectral region. This finding means that the binuclear center of oxygen reduction keeps a very similar structure and holds the same ligands in the studied conditions. The further investigation in search of differences should be performed in the 4000-2000 cm^-1^ IR region where water ligands absorb.

**Abbreviations:**

A: ferrous-oxy intermediate
ATR: Attenuated total reflectance
BG: background
BNC: binuclear center
C*c*O: Cytochrome *c* oxidase
E: one-electron reduced intermediate
F: ferryl intermediate
FRCO: fully-reduced CO-inhibited enzyme
FTIR: Fourier transform infrared
IR: infrared
NHE: Normal Hydrogen Electrode
N-side: negatively charged side of the membrane
P_R/M_: “peroxy”
intermediates; P-side: positively charged side of the membrane
R: fully-reduced intermediate
TR: time-resolved
WT: wild type

## 1. Introduction

Cytochrome *c* oxidase is the terminal complex of the respiratory chain of mitochondria, and some aerobic bacteria and archaea. C*c*O possesses four redox centers, in the direction of electron flow: Cu_A_, heme *a*, heme *a*_*3*_, and Cu_B_. The last two centers form the binuclear center (BNC) where catalysis takes place. C*c*O receives electrons from a small soluble enzyme—Cytochrome *c*—from the positively charged side of the membrane (P-side). Cytochrome *c* passes electrons to Cu_A_ one by one. For oxygen reduction catalysis, the BNC of C*c*O must receive four protons (“chemical” protons; because they go for the chemistry of the catalytic cycle) from negatively charged side of membrane (N-side). This reaction is very exergonic, and the energy released is used to translocate four more protons across the membrane (these are called “pumped” protons) (1–4). Two H-conductive channels serve to transfer the protons needed for the chemistry and pumping: D- and K-channels, named after the conserved residues D-124 and K-354 (here *Paracoccus denitrificans* numbering used because this work was performed on *P. denitrificans* C*c*O enzyme), respectively (5–11). The catalytic cycle includes six relatively stable at room temperature intermediates determined mainly by time-resolved visible and Raman spectroscopies: **R** (fully-reduced) → **A** (ferrous-oxy) → **P**_**R/M**_ (“peroxy”; **P**_**R**_ formed from fully reduced and **P**_**M**_ formed from two electron reduced enzyme, electrons are located at the BNC) → **F** (ferryl) → **O/O**_**H**_ (fully-oxidized “resting”/”pulsed”) → **E/E**_**H**_ (one-electron reduced “resting”/”pulsed” states). (**Fig.1**, reproduced from (12)).

The catalytic cycle can be speculatively divided into two halves: oxidative (**R→O**) and reductive (**O→R**) halves (Fig. 1). This work focuses on the fully-oxidized intermediate that is measured from both sides of the catalytic circle in experiments. In this intermediate all four redox centers are present in a fully-oxidized state. The fully-oxidized intermediate may exist in different forms with different rates of reduction and reactivity towards external ligands (13–14). The “slow” form has a maximum in the Soret region below 418 nm and is characterized by the slow kinetic of cyanide binding to the oxidized heme *a*_*3*_. The absorption maximum of the “fast” form red-shifted by several nm. The enzyme in the “slow” form can be obtained when the protein is incubated or isolated at a low pH value. When C*c*O is extracted from the membrane, it exists in the slow or “resting” **O** form. Moreover, this form is heterogenic and is thought of as simply an artefact of preparation method. Incubation of “fast” oxidase with chloride or bromide at low pH produces forms of the enzyme that are similar to the “slow” form of the enzyme (13). Reduction of the enzyme in **O** state does not couple to proton pumping (15–17). “Slow” oxidase shows characteristic EPR signal arising from the BNC, at g ≈ 12 and g = 2.95, that are not seen in the fast form of enzyme. Reduction of this enzyme and its re-oxidation produces a fast or “pulsed” **O**_**H**_ state. The reduction of this intermediate is accompanied by the pumping of two protons across the membrane (15,18). It was predicted that **O**_**H**_ state relaxes to the **O** state within about ∼ 30 s at room temperature (18). It was thought that the “resting” and “pulsed” states differ in their redox properties or ligands of the BNC (18). Later it was proposed that the difference between these intermediates are in the presence of water molecules or their reorientation in the BNC or in close proximity of it (19,20). Later, Jancura *et al.* characterized **O** and **O**_**H**_ protein states by visible kinetic and EPR spectroscopies and found no difference between these two states. They concluded that the reduction potentials and the ligation states of heme *a*_*3*_ and Cu_B_ are the same for C*c*O in the **O** and **O**_**H**_ states. Nowadays, the structure of the BNC in **O** and **O**_**H**_ states are supposed to be as follows: Fe^III^-OH-Cu^II^-HOH Tyr(280)-O^-^ (as in Fig. 1).

**Fig. 1.**
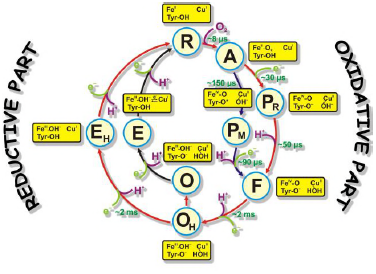
The catalytic cycle of C*c*O with structures of resolved intermediates. Protons taken by the BNC for the reaction chemistry are shown in magenta. Electrons taken by the BNC are marked in light green. Time constants of the appearance of reaction intermediates are also shown in green. The proposed structures of the catalytic site represent only the BNC structure and provide no information about the other two redox centers (Cu_A_ and heme *a*).

This paper presents a study of **O**→**R** transition on statically-resolved FTIR redox spectra of wild type (WT) enzyme at pH 6.0 and 9.0, and **R**→**O** transition of WT at two pH values (6.0 and 9.0); on D124N variant at pH 9.0; and N131V mutant at pH 6.5 and 9.0. **R** intermediate is assumed to be the same in all studied cases and only oxidized intermediates may vary. Here, static and time-resolved FTIR spectroscopy demonstrates that there are no clear differences in the oxidized intermediate irrespective of type of the mutant enzyme or WT and the way the **O/O**_**H**_ intermediate is produced and measured, i.e. there is no difference between “resting” and “pulsed” fully-oxidized intermediate within the measured IR spectral range.

## 2. Materials and methods

### 2.1 Sample preparation

For the preparation of “ATR-ready” sample, the variants of the enzyme were depleted of the detergent as originally described in (21) with some modifications (22–24). The “ATR-ready” sample of C*c*O was placed on a microprism slightly polished by aluminum powder ATR (Attenuated total reflectance) microprism (Sens IR Technologies (Danbury, USA), three-bounce version, surface diameter 3 mm). The sample was dried under soft N_2_ flow or under a tungsten fiber lamp positioned at a distance of ∼30 cm from the sample film, then, rewetted with “working” buffer (= buffer in which the experiments are undertaken). In all experiments, the enzyme film was stabilized overnight, then covered with a specially designed chamber either for equilibrium FTIR measurements (22) or for a time-resolved approach (24) (also see two sections below). Sample preparation for the static experiment included pumping of “working buffer” overnight in order to stabilize the protein film. In the morning, such a sample was ready for static experiments. Sample preparation for the kinetic FTIR **R→O** measurements also included overnight incubation but without pumping and **FRCO** (Fully-reduced CO-inhibited) complex preparation with buffer contained 100% CO gas. Formation of **FRCO** complex was followed by the appearance of heme *a*_*3*_-C≡O band at 1965 cm^-1^ and took approximately 40 to 80 minutes.

### 2.2 Equilibrium O→R FTIR spectra

Equilibrium redox titration setup built in Helsinki Bioenergetics Group (Ref. (22) and (12), **Figs. 9, 10** therein) allows to measure the reductive half of the catalytic cycle (Fig. 1). Equilibrium redox FTIR spectra have been measured in a broad pH region before (25,26). The buffer composition was the following: 400 mM of K_2_SO_4_, 25 mM of potassium acetate, 25 mM of potassium phosphate, and 25 mM of boric acid. The spectra were produced electrochemically with a homemade flow-by electrochemical cell (27). The picture of the cell with its description can be found in (22) and (27). In short, gold grains were used as the working electrode, a platinum-plated titanium grid—as the counter electrode and Ag/AgCl/saturated KCl—as the reference electrode. The “working buffer” with mediators was continuously pumped through the working electrode to the ATR chamber. The cell was connected to a Princeton Applied Research potentiostat (Oak Ridge, Tennessee, USA). To equilibrate the enzyme film with the potential of the working electrode, two mediators were used: 1,2-diaminocyclohexane-N,N,N’,N’-tetraacetic acid + Fe, midpoint potential +95 mV; and ferrocene acetate, +370 mV. The kinetics of equilibration of the enzyme film with the potential of the working electrode was controlled in the visible region spectrophotometrically. The **O→R** spectra measurements were performed as follows. First, the enzyme film was equilibrated at acidic pH at -300 mV (vs NHE, Normal Hydrogen Electrode) for 15 min. Then, a background (BG) spectrum in the infrared (IR) region was measured. Then, the potential was set at +480 mV, again allowing 15 min for equilibration. At this point, a spectrum in the IR region was taken. Immediately after that, BG in the IR region was collected. The potential was then switched back to -300 mV and the procedure was repeated many times to collect FTIR spectra with good signal-to-noise ratio. Then, the pH of the “working buffer” was changed to another value (in this case to pH 9.0) and a series of **R→O** and **O→R** spectra were measured and summed up.

To follow oxidoreduction, an FTIR spectrometer Bruker IFS 66/s (Ettlingen, Germany) equipped with a fast MCT detector and a visible spectrophotometer USB2000 (Ocean Optics) (Largo, USA) were used. FTIR spectra were measured in the spectral range 4000-500 cm^-1^ with 4 cm^-1^ spectral resolution and IR scanner frequency 80 kHz. The number of scans in BG and sample spectra was 2000; apodization function, Blackman-Harris 3-Term. The control of the status of oxidoreduction was achieved by visible spectroscopy at 445 nm. The light guide from a visible spectrophotometer was connected to the ATR-chamber. The data acquisition system described above produced one **R→O** and one **O→R** spectrum that were summed up. All measurements were performed at room temperature. For more detailed description of the procedure of spectra collection see Ref (25).

The whole data acquisition process—initializing, sending commands to/from Ocean Optics and FTIR Opus software, and collecting data—was controlled by *Titrator* software designed in the Helsinki Bioenergetics Group by Dr. Nikolay Belevich.

### 2.3. Time-resolved R→O FTIR spectra

With help of time-resolved **R→O** FTIR spectra, the oxidative part of the catalytic cycle of C*c*O was investigated. Time-resolved **R→O** FTIR spectra were measured in the following conditions: WT at pH 6.0 and 9.0; N131V at pH 6.5 and 9.0; and D124N at pH 9.0. These spectra were measured in order to distinguish similarities and dissimilarities between **O** and **O**_**H**_ states in different conditions as much as possible. To perform time-resolved measurements of the **R→O** transition, a special setup was constructed. The core of this setup is an FTIR spectrometer Bruker IFS 66/s equipped with a fast MCT detector and an ATR-cell. The ATR-cell is covered with a special chamber for time-resolved oxygen reaction measurements. Enzyme on ATR-prism is prepared in **FRCO** form. CO is photolyzed from C*c*O **FRCO** enzyme by a laser that is delivered by a laser guide connected to the chamber. Oxygenated buffer is injected very close to the enzyme film. A light guide from a visible spectrophotometer is constructed in the ATR-chamber as well as an inlet and outlet of the pump. For more details and for pictures see Ref. (24) and Fig. 1 therein and Ref. (12) **Figs. 12-14** therein.

The “working buffer” for time-resolved FTIR measurements included 100 mM glucose (Merck (Darmstadt, Germany)), 260 µg/mL catalase (Sigma-Aldrich (St. Louis, Missouri, United States)), 3.3 mM ascorbate (Prolabo (Leicestershire, UK)), 10-100 µM hexaaminruthenium (III) chloride (Sigma-Aldrich (St. Louis, Missouri, United States)) (the amount of the last component depended on C*c*O variant), and 670 µg/mL glucose oxidase (Roche (Basel, Switzerland)).

Before each experiment, the sample quality control was conducted (24). Four procedures were applied in order to test the stability and functionality of the BNC of C*c*O. First, the intensity and the shape of the band at 1965 cm^-1^ was analyzed. Second, CO dissociation from heme *a*_*3*_ was followed by visible time-resolved spectroscopy with 1 ms temporal resolution. Third, CO dissociation in the dark from heme *a*_*3*_ was followed by time-resolved (TR) rapid-scan FTIR in the region around the band at 1965 cm^-1^. Finally, CO dissociation from **FRCO** compound at 1965 cm^-1^ was measured after oxygenated buffer addition, delay of 3 s, and laser flash. This last test was performed before, in the middle, and at the end of each experiment. It showed the amount of enzyme capable of performing the oxygen reaction.

**FRCO→O** spectra were acquired as initially described in Ref. (24). Spectra were measured in 1850-950 cm^-1^ region, which was cut out by an interference filter. The spectra were collected in the rapid-scan mode with ∼ 46 ms temporal resolution and 8 cm^-1^ spectral resolution. Together with oxygen injection, rapid-scan acquisition started and was followed by a 3 s delay and the laser flash. The flow pump was stopped during the kinetic data collection and switched on immediately after to speed up the C*c*O re-reduction. This cycle was repeated many times to get spectra of a good signal-to-noise-ratio. A global fitting procedure in *Matlab* software (Natick, MA, USA) was applied to extract the slow part of kinetical spectra. The fast part of kinetical spectra was calculated as a difference between the spectrum obtained by the average of several time points before the laser flash and the spectrum obtained immediately after it. The sum of the fast and the slow components gave resulting kinetic spectrum in case of mutated C*c*O. The kinetic spectrum in case of WT enzyme was calculated as a spectral jump after the laser flash because oxygen reaction in WT enzyme proceeds faster than temporal resolution of the measurement setup.

**FRCO** photolysis was measured separately at pH 6.5 on WT variant and subtracted from each **FRCO→O** spectra to produce **R→O** spectra presented in the figures in the section **“Results”**. It was shown in (28) that **FRCO** photolysis on WT does not depend on pH values.

All measurements of kinetic **R→O** transitions and **FRCO** photolysis were performed in ice-cooled conditions in order to slow-down the reaction rate and increase concentration of oxygen until reaching 2.4 mM. For more details see Ref. (20,24).

### 2.4 Data analysis

All data analysis and figure preparations were performed in *Matlab* software.

## 3. Results and discussions

Here, the IR spectra are compared in acidic conditions on static and time-resolved redox transitions (**O→R** for static transitions and **R→O** for the time-resolved). Altogether 6000 FTIR **O→R** static spectra were collected at pH 6.0 on WT (Fig. 2, red spectrum), 173 kinetic spectra of WT at pH 6.0 (**Fig.2**, dark green spectrum), and 338 kinetic spectra of N131V at pH 6.5 (Fig. 2, dark blue spectrum).

**Fig. 2.**
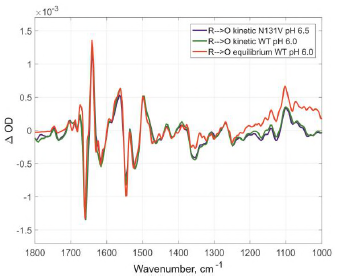
Comparison of the R→O transitions in three different conditions. Time-resolved mode for N131V at pH 6.5 in dark blue and for WT at pH 6.0 in dark green; equilibrium mode for WT at pH 6.0 in red. All spectra normalized to 1 mM (kinetic – by amplitude of 1965 cm^-1^ band, equilibrium – by 1661/1641 peaks difference).

Small differences between static and kinetic spectra are due to difference in their spectral resolution which is 4 cm^-1^ for static and 8 cm^-1^ for the kinetics spectra.

Similar results were observed under alkaline conditions: 296 kinetic spectra for D124N at pH 9.0 (Fig. 3, dark blue), 218 for N131V 9.0 (sFig. 3, dark green), 82 for WT (Fig. 3, red), and 6000 co-additions for WT in static conditions (Fig. 3, light-green).

**Fig. 3.**
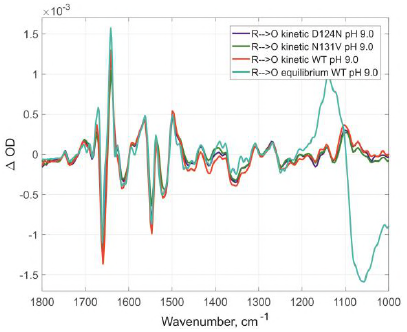
Comparison of R→O transitions in four different conditions at pH 9.0. Time-resolved spectra for D124N (in dark blue), N131V (in dark green), and WT (in red); equilibrium spectrum for WT (light green). All spectra were measured at pH 9.0 and normalized to 1 mM (kinetic – by amplitude of 1965 cm^-1^ band, equilibrium – by 1661/1641 peaks difference).

No differences were found between oxidized intermediates in the IR region 1800-1000 cm^-1^ for all studied cases in both acidic and alkaline conditions. There are no visible differences between static and time-resolved **R→O** FTIR spectra of WT and mutated enzymes in the IR region 1800-1000 cm^-1^ that prove that the BNC of these variants and in these pH conditions are very similar. It does not matter what side we go **R→O** or **O→R**, oxidized intermediate provides the same findings and no difference between **O** and **O**_**H**_ is observed in the 1800-1000 cm^-1^ infrared window for all conditions studied. Further, the difference should be sought in the so-called “water region.”

## 4. Conclusions

This paper compared the oxidized intermediates derived from different variants (WT, D124N, and N131V) and using different approaches (static **O→R** spectra acquisition and time-resolved **R→O** spectra). To perform this work, two special setups were constructed: one for static measurements and one for time-resolved measurements. This work provides an overview of the oxidized intermediates prepared under very different conditions: different mutants and pH values. No differences are evident between all of the conditions tested, except that for static spectra spectral resolution was set at 4 cm^-1^ and for time-resolved at 8 cm^-1^. This similarity in oxidized intermediates under different conditions means that the structures of BNC in all these cases (**O** and **O**_**H**_ states) are very similar from the FTIR fingerprint point of view. Further investigation in the IR region 4000-2000 cm^-1^ is required to search for differences between various oxidized intermediates.

## Author contributions

E.G. designed research; E.G. performed research; E.G. and R.K. analyzed the data; E.G. wrote the manuscript.

## Acknowledgements

EG was supported by Center of International mobility (CIMO) and Informational and Structural Biology (ISB) Graduate School. The work was supported by the Sigrid Juselius foundation, Biocentrum Helsinki and the Academy of Finland (project numbers 200726, 44895, and 115108). I warmly thank Dr. Nikolai Belevich for an excellent technical assistance, Dr. Timur Nikitin for critical reading of the manuscript and Jennifer Rowland for language correction.

